# Molecular structures enumeration and virtual screening in the chemical space with RetroPath2.0

**DOI:** 10.1101/158147

**Authors:** Mathilde Koch, Thomas Duigou, Pablo Carbonell, Jean-Loup Faulon

## Abstract

**Background:** Network generation tools coupled with chemical reaction rules have been mainly developed for synthesis planning and more recently for metabolic engineering. Using the same core algorithm, these tools apply a set of rules to a source set of compounds, stopping when a sink set of compounds has been produced. When using the appropriate sink, source and rules, this core algorithm can be used for a variety of applications beyond those it has been developed for.

**Results:** Here, we showcase the use of the open source workflow RetroPath2.0. First, we mathematically prove that we can generate all structural isomers of a molecule using a reduced set of reaction rules. We then use this enumeration strategy to screen the chemical space around a set of monomers and predict their glass transition temperatures, as well as around aminoglycosides to search structures maximizing antibacterial activity. We also perform a screening around aminoglycosides with enzymatic reaction rules to ensure biosynthetic accessibility. We finally use our workflow on an *E. coli* model to complete *E. coli* metabolome, with novel molecules generated using promiscuous enzymatic reaction rules. These novel molecules are searched on the MS spectra of an *E. coli* cell lysate interfacing our workflow with OpenMS through the KNIME analytics platform.

**Conclusion:** We provide an easy to use and modify, modular, and open-source workflow. We demonstrate its versatility through a variety of use cases including, molecular structure enumeration, virtual screening in the chemical space, and metabolome completion. Because it is open source and freely available on MyExperiment.org, workflow community contributions should likely expand further the features of the tool, even beyond the use cases presented in the paper.

## 1. Introduction

The number of known chemical reactions is huge, at the time this manuscript was written there were ~84 million single- and multi-step reactions in the Chemical Abstract Service database (CAS) [1]. Yet, many reactions in CAS are redundant because the same reactions are applied to different reactants. Identifying identical reactions can be performed by computing reaction rules. Reaction rules represent reactions at the reaction center only. In other words, a reaction rule comprises only the substructures of the reactants and the products for which the atoms are either directly involved in bond rearrangements or are deemed to be essential for the reactivity of the reaction center. While a set of reaction rules is of course not available for all known chemical reactions, rules have been compiled for focused applications, such as retrosynthesis planning [2, 3], the discovery of novel chemical entities in medicinal chemistry [4], xenobiotic (including drug) degradation [5], metabolomics [6], and metabolic engineering [7-10]. Depending on the application, the number of rules varies from less than one hundred to few thousands, but in all cases the number of known reactions per application far exceeds the number of rules (there are for instance more than 14,000 reactions in metabolic databases such as MetaNetX [11]). There are several ways of coding reaction rules (for instance, BE-matrices [12] and fingerprints [13]) but most of the time the rules can be represented by reaction SMARTS [14], as it is done in the current paper.

The purpose of reactions rules is to generate reaction networks. The rules can be used in a forward manner to find for instance the metabolic degradation products of a drug, or in a reverse manner to find the reactions producing a desired product from a set of available reactants. In this later usage one produces retrosynthesis reaction networks. Several tools have been developed in the past to generate (retrosynthesis) reaction networks and reviews are available for synthesis planning [2, 3] and for metabolic engineering [15]. Disregarding if the rules are applied in a forward or reversed manner, network generation tools are making use of the same core algorithm. Starting from a source set of compounds the core algorithm applies the rules in an iterative fashion either a predefined number of times or until a sink set of compounds have been produced. At each iteration, the algorithm fires the rules on the source set producing *new* molecular structures and determines the *new* source set of molecules the rules will be fired upon at the next iteration. That set must comprise molecules that have not been processed before. Further details on the core algorithm and the differences between the various implementations are provided in Faulon *et al.* [16] and Delépine et al. [17].

In the current paper we make use of an open source workflow (RetroPath2.0 [17]), which follows the above core algorithm. This workflow is not based on original codes but instead was constructed entirely by assembling KNIME nodes [18] developed by the cheminformatics community (primarily RDKit nodes [19]). RetroPath2.0 is the first open source release of a retrosynthesis reaction network generation, its usage in the current paper beyond network generation demonstrates its versatility.

*As already mentioned, reaction network generation tools coupled with reactions rules have been developed and used primarily for synthesis planning and metabolic engineering, but can they be used to enumerate molecules (isomers for instance) and more generally to search chemical structures in the chemical space?*

In principle yes if one can devise reaction rules enabling the production of any molecule in the chemical space. Such a set of rules necessarily exists for all know molecules (such as those in the CAS database) since they have been produced through either natural or synthetic chemical reactions. In practice and as already stated, reaction rules so far developed are application limited. Yet, within their respective application fields, specific rules have been used to discover novel molecules and reaction pathways. Taking experimentally validated examples, the rules associated with the ligand-based de novo design software DOGS (inSili.com LLC) [4] have enabled the production of new chemical entities inhibitors of DAPK3 (death-associated protein kinase 3) [20], metabolic rules for promiscuous enzymes have allowed the discovery of novel metabolites in *E. coli* [21] and have also been used to engineer metabolic pathways producing 1,4-butanediol [9] and flavonoids [22].

Going beyond application limited reaction rules, the main contribution of the present paper is to propose a set of transformation rules that enables the generation of any isomer of any given molecule of the chemical space. Precisely, we prove the claim that any isomer of any given molecule of *N* atoms, can be reached applying at most O(*N*^2^) rules.

As illustrations, our transformation rules are used to screen the chemical space for structures that are similar to a given set of well-known monomers and to search aminoglycosides structures maximizing antibacterial activities. The compounds produced by our rules are not necessarily chemically accessible, since our transformation rules are not constructed based on chemical synthesis schema. To probe the (bio)synthetic accessibility of our solutions, we also perform search in the (bio)chemical space using enzymatic reaction rules. The enzymatic rules are also used to propose novel molecules completing *E. coli* metabolic network and for which masses are found in cell lysate mass spectra.

All results presented in this paper have been produced making use of the open source workflow RetroPath2.0. RetroPath2.0 and the associated data are provided as Supplementary and can be downloaded at MyExperiment.org. The only differences between the various usages we have made of the RetroPath2.0 are within 1) the set of reaction rules and 2) the way molecules are selected at each iteration during the network generation process.

## 2. Results and Discussions

The purpose of this section is to showcase the versatility of the RetroPath2.0 by taking use cases of interest to the community. We first propose reaction rules to enumerate isomers (2.1), we then use the rules to screen in the chemical space structures that are similar to some known monomers (2.2) and compute property distribution (Glass transition temperature) in both the Chemical Space and PubChem, we next use a QSAR to search aminoglycosides types molecules for which antibacterial activity is maximized using both isomer transformation rules and enzymatic rules (2.3), and we finally use enzymatic rules to find novel metabolites in *E. coli* and annotate the MS spectra of an *E. coli* cell lysate interfacing RetroPath2.0 with OpenMS [23] (2.4).

### 2.1. Isomer enumeration

Isomer enumeration is a long-standing problem that is still under scrutiny [24, 25]. Our intent here is not to provide the fastest enumeration algorithm but to demonstrate how RetroPath2.0 can perform that job once appropriate reaction rules are provided. However, we provide in supplementary Figure S1 a comparison of Retropath2.0’s execution time with the OMG and PMG software tools [25,26] specifically dedicated to isomers enumeration. Retropath2.0 is found faster than OMG but slower than PMG. Thereafter, we outline below two approaches making use of RetroPath2.0. The first is based on the classical *canonical augmentation* algorithm [27] and the second consists of iteratively transforming a given molecule such that all its isomers are produced. We name this latter approach *isomer transformation.* In both cases we limit ourselves to structural (constitutional) isomers, as there already exist workflows to enumerate stereoisomers [28].

*Canonical augmentation.* The principle of canonical augmentation, which is an orderly enumeration algorithm, is to grow a molecular graph by adding one atom at a time and retaining only canonical graphs for the next iteration [27]. The algorithm first proposed by Brendan McKay has been used to generate the GDB-17 database of small molecules [29]. The original algorithm has also been modified such that at each step a bond (not an atom) is added to the growing molecules [25]. In the present implementation we use the original McKay algorithm [27], consequently, the number of iteration is the number of atoms one wishes the molecule to have. The algorithm can easily be implemented into RetroPath2.0 by choosing as a source set a single unbonded atom, and a rule set depicting all possible ways an atom can be added to a molecular graph (see method section for more information). Considering that an atom can be added to a growing molecule through one, two, or more bonds (depending on its valence), the set of reaction rules is straightforward however cumbersome if one starts to consider all possible atoms types. For this reason we limit ourselves to carbon skeleton as it is usually done in the first step of isomer enumeration algorithm. Figure 1 below depicts the set of rules that generate all triangle free carbon skeletons.

**Figure 1.**
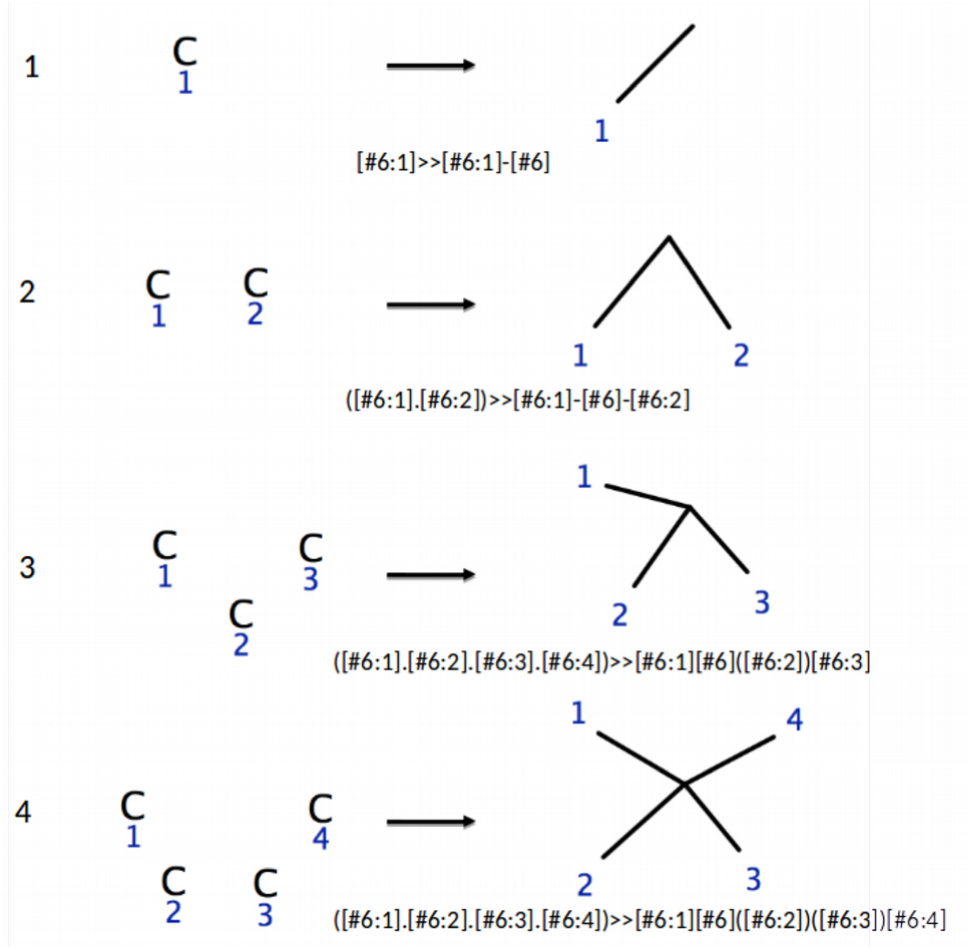
Reaction rules for canonical augmentation of carbon skeletons. The corresponding reaction SMARTS string is provided for each rule.

We note that rules R_2_ to R_4_ will generate cycles since the added atom is attached to the growing molecule by 2 to 4 four bonds, thus only rule R_1_ is necessary to grow acyclic molecules (alkanes for instance). The Table below provides the numbers of alkane structural isomers found up to 18 carbon atoms running RetroPath2.0 with rule number 1 in Figure 1.

The numbers agree with earlier calculations [30]. For a given number of carbon atoms (*N*), the canonical augmentation generate all alkanes from 1 to *N* carbon atoms, while the isomer transforming enumeration generate alkanes having only *N* carbon atoms, one can thus verify that at any given number of carbon atoms *N*, the numbers of structures generated by the canonical augmentation algorithm equals the sum of numbers of isomers generated by the transformation algorithm up to *N*.

*Isomer transformation.* The isomer canonical augmentation algorithm becomes more complex when one starts to consider different atom and bond types. To overcome these difficulties the idea of the transformation enumeration approach is to start with one fully-grown molecule to which one applies all possible transformations such that all the structural isomers of the initial molecule are generated. This approach can be implemented in RetroPath2.0 using a hydrogen saturated molecule as a source and a reaction rule set enabling to transform the molecule while keeping the correct valence for each atom. Because atom valences are maintained the total number of bonds must remain the same after the transformations have taken place. In order to maintain the number of bonds constant, for any reaction rule the number of bonds created must equal the number of bonds deleted.

RetroPath2.0 applies a reaction rule to a given molecule by first searching all occurrences in the molecule of the subgraph representing the reactant (left side of the rule). To this end the labels on the subgraph are removed. Then for each occurrence of the unlabeled subgraph in the molecule, the labels are restored and the bonding patterns on the molecule are changed accordingly. The process is illustrated in the Figure 2 below where it can be seen that rules *R*_a_ and *R*_b_ are identical (i.e. they produce the same solutions). In general, two rules *R*_a_=(L_a_, A) and *R*_b_=(L_b_,B) will produce the same solutions if a one-to-one mapping π can be found between the labels L_a_ and L_b_ of the rules such that the set of edges (A) in *R*_a_ is transformed by π into the edges (B) of *R*_b_, i.e. π(A)=B.

**Figure 2.**
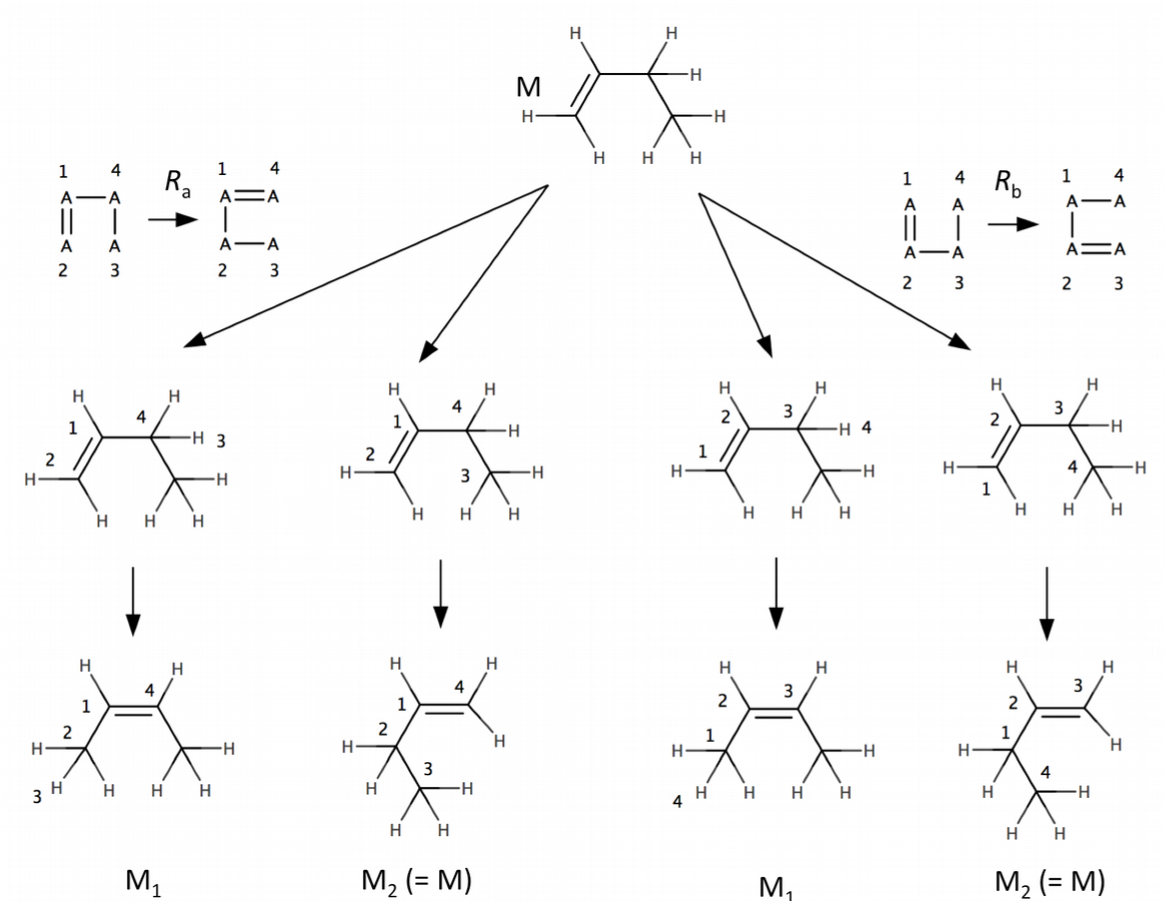
Identical rules. There are two different ways (two different possible matchings for the reactants of the rules) of applying rules *R*_a_ and *R*_b_, each rule produces molecules M_1_ and M_2_. The molecules produced by *R*_a_ are identical to those produced by *R*_b_ because the rules are identical. *R*_a_ is identical to *R*_b_ because when applying the one-to-one label mapping π(1,2,3,4)= 2,1,4,3 on the edges of the R_a_ one obtains the edges of *R*_b_.

#### Claim

The 19 rules described in Figure 3 allow us to generate all isomers of a given molecule at most 3/4*(*N*^2^ – *N*) iterations, where *N* is the number of atoms, respecting the following constraints: the maximal valence is 4 and there cannot be two double bonds on the same atom in a 3 or 4 membered ring.

**Figure 3.**
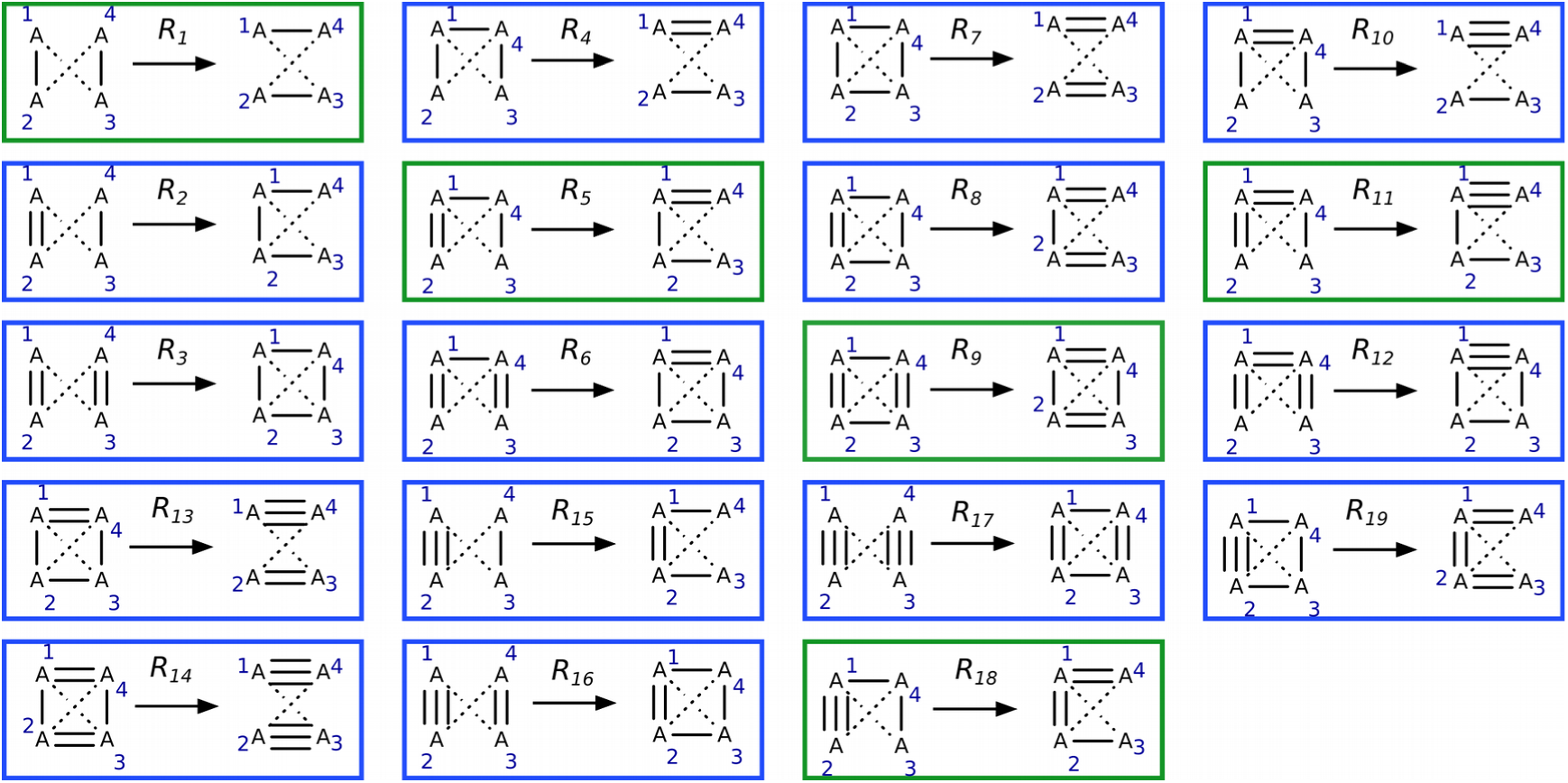
Isomer transformation rule set. All reactions rules are solutions of system of equations (2) and are non identical (see text and Figure 2 for definition of identical rules). Reactions in green move bonds around without creating or deleting cycles. Reactions in blue change bond order by creating or deleting at least one cycle. To each reaction corresponds a reverse reaction. The reverse reaction of *R*_1_ is *R*_1_, for *R*_2_ it is *R*_4_, for *R*_3_ : *R*_7_, for *R*_5_: *R*_5_, for *R*_6_ : *R*_8_ and the reverse reaction of *R*_9_ is *R*_9_. The reverse reaction for *R*_10_ is *R*_15_, for *R*_11_, *R*_18_, for *R*_12_, *R*_19_, for *R*_13_, *R*_16_ and for *R*_14_, *R*_17_. The bond order a_13_ and a_24_ can take any value from 0 to 3. The full list of rules excluding triple bonds can be found in Figure S2.

##### Lemma 1

The minimal number of bonds one can change is 4 and the 19 rules described in Figure 3 generate all minimal transformations respecting the following constraints: the maximal valence is 4 and there cannot be two double bonds on the same atom in a 3 or 4 membered ring.

##### Proof

The minimal transformation one can perform consists of deleting one bond and creating another one. Since the bond created must be different than the one deleted at least three atoms (A_1_, A_2_, A_3_) must be involved. Let a_12_, a_13_, and a_23_ be the bond order between the three atoms and let b_12_, b_13_ and b_23_ the bond order after the reaction has taken place. Because the atom valence is maintained the following system of equations holds:

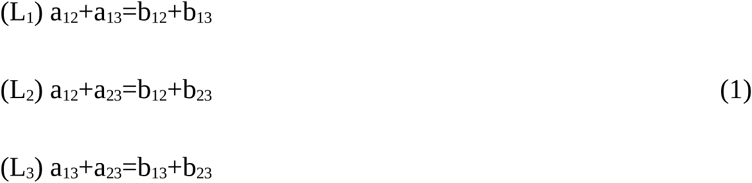

(L_1_)+ (L_2_) – (L_3_) -> a_12_ = b_12_, which implies a_23_ = b_23_ and a_13_ = b_13_.

It is therefore impossible to proceed to a minimal transformation with only 3 bonds involved.

Let us consider 4 atoms. There are 6 possible bonds between those atoms. Let us consider that we are changing 4 bonds, since we aim to find minimal transformations. Let us call a_13_ and a_24_ the two fixed bonds, without loss of generality.

Valence conservation (with b_13_ = a_13_ and b_24_ = a_24_) gives us the following system:

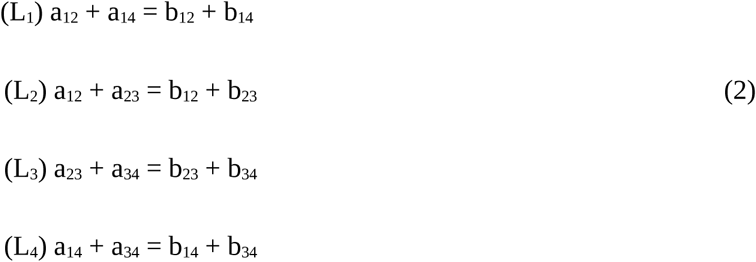

We can notice that (L_1_) + (L_3_) = (L_2_) + (L_4_): we therefore have a system of 3 equations with 4 unknowns, so we can set an unknown and calculate the other solutions.

As we are looking for minimal transformations, we can assume that we are changing a bond order by 1 on this solution we can set. Since valence is conserved, if a bond order is increased, then a bond order from the same atom has to be decreased. As the problem is perfectly symmetrical in all variables at this point, and we can us assume without loss of generality (at least one bond has to be deleted) that b_12_= a_12_ -1. Then solving the system immediately gives us b_14_ = a_14_ + 1, b_23_ = a_23_ +1 and b_34_ = a_34_ -1. This system can only be solved in our case (positive bond orders, no quadruple bonds) if a_14_ and a_23_ are either 0, 1 or 2 and a_12_ and a_34_ are either 1, 2 or 3. This means that we have at most 81 (3^4^) cases for initial bond orders where our isomer problem has a solution. However, this solution space can be further reduced by problem symmetry arguments. We can see that the roles of a_12_ and a_34_ are symmetrical, as well as the roles of a_23_ and a_14_.

Let us call A_1_ the atom with the highest considered sum of bound orders (neglecting the fixed orders a_13_ and a_24_). Therefore, it is such that

a_12_ + a_14_ >= a_12_ + a_23_ (higher sum of bound orders than A_2_), or a_14_ >= a_23_ (Condition 1)

a_12_ + a_14_ >= a_14_ + a_34_ (higher sum of bound orders than A_4_), or a_12_ >= a_34_ (Condition 2)

a_12_ + a_14_ >= a_23_ + a_34_ (higher sum of bound orders than A_3_), is automatically verified when the other 2 are verified.

Condition 1 is not respected when a_23_ = 2 and a_14_ = 0 or 1 or when a_23_ = 1 and a_14_ =0, without constraints on a_12_ and a_34_: (2+1) * 9 = 27 solutions. For the same reason, 27 solutions do not respect condition 2. The solutions that do not respect both condition 1 and condition 2 are (2+1) * (2+1) = 9. By symmetry arguments, we therefore reduce the solution space from 81 to 81 – 27 – 27 + 9= 36. These 36 reaction rules are presented in Figure 4.

We can further reduce the solution space by considering that the maximum atom valence is 4. The solutions that do not respect this rule are such that a_12_ + a_14_ = 5, so a_12_ = 3 and a_14_ =2 (and this automatically verifies conditions 1 and 2). Since there are no constraints on a23 and a34, we have 9 such solutions: the solution space has been reduced to 36 – 9=27 reactions.

One more constraint, imposed by 3D conformation of the molecule, is that there cannot be two double bonds on the same atom in a 3 or 4 membered ring.

This must be true for our initial molecule as well as for the produced molecule. For the initial molecule (as can be seen in the 4 black rules under rule 13 in Figure 4), when a_12_ = a_14_ = 2, since a_34_ >1 (bond whose order will be reduced), there is a cycle if a_23_ ≠ 0. There are therefore 4 solutions where the initial molecule is invalid: when a_12_ = a_14_ = 2, and a34 is 1 or 2 (smaller than a_12_) and a_23_ is 1 or 2.

This must also be true for the produced molecule. Two double bonds will be produced around atom 1 with a_12_ = 3 and a_14_=1 (this can be seen in the 4 black rules under rule 18 in Figure 4). There will be a cycle if a_34_ ≠ 1. There are therefore 4 solutions where the produced molecule is invalid: when a_12_ = 3, a_14_=1,and a_34_ is 2 or 3 and a_23_ is 0 or 1 (smaller than a_14_).,

Since these solutions respect valence and problem symmetry, they are not included in the previous solution space reductions and therefore the solution space is reduced to 27 – 8 =19 solutions. A summary table of solution space reduction is given in Table S1. Since we have found 19 different working solutions for all the cases we have left, we have proved that the minimal number of bonds one can change is 4 and the 19 rules described in Figure 3 generate all these minimal transformations.

**Figure 4.**
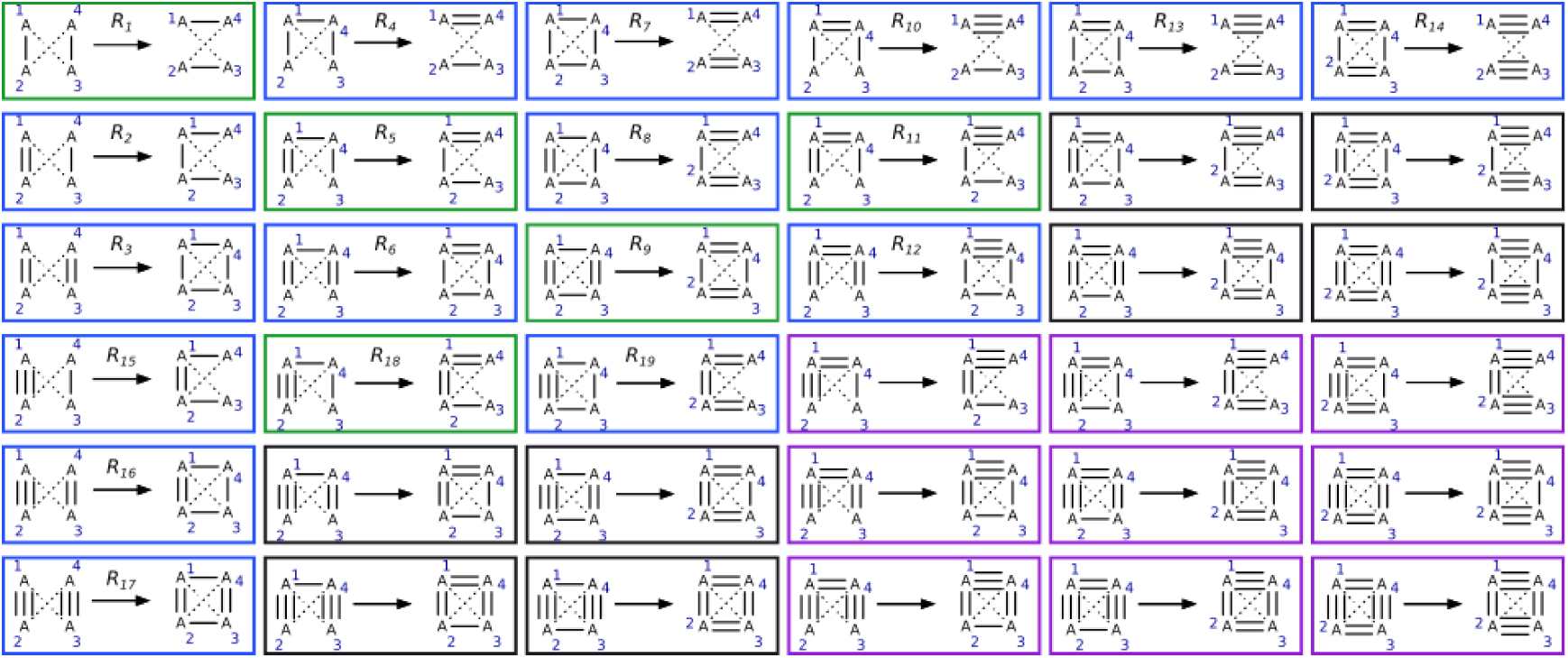
Rules before solution space reduction due to valence and structure considerations. Reactions in green move bonds around without creating or deleting cycles. Reactions in blue change bond order by creating or deleting at least one cycle. Reactions in purple were deleted because valence is limited to 4, and reactions in black were deleted because there cannot be two double bonds on the same atom in a 3 or 4 membered ring.

##### Lemma 2

Let us consider M_b_ an isomer of M_a_. We can apply a rule from this set of 19 rules that will reduce the sum of absolute order differences between those two molecules by at least 2 and at most 4.

##### Proof

Let (a_ij_) be the order of bonds in M_a_, (b_ij_), j Є [2,*N*], i Є[1,j-1], the order of bonds in M_b_, where *N* is the number of atoms in M_a_ and M_b_. Since M_b_ is different of M_a_, we can find i,j such that a_ij_ > b_ij_. By valence conservation in atom A_j_, we can find k such that a_jk_ < b_jk_, and by valence conservation of atom A_k_, we can also find l such that a_kl_ > b_kl_. Therefore, we are considering 4 atoms and 4 bonds between those atoms, with at least 3 of their orders changing by 1. According to Lemma 1 the minimal number of bonds one can change is 4, so we will also have to change the bond order between A_i_ and A_l_. We are therefore considering a minimal transformation, so we know thanks to Lemma 1 that we can apply a rule from our set of rules to generate that transformation. Let us call M_a’_ the molecule produced that way, and a_ij’_ its bond orders. Let us now calculate the sum of orders of M_a’_. Then, by applying the rule, we have a_ij’_ = a_ij_ -1 and therefore ∣b_ij_ – a_ij’_∣ = ∣b_ij_ – a_ij_∣ - 1. For the same reason, ∣b_kl_ – a_kl’_∣ = ∣b_kl_ − a_kl_,∣ - 1. Moreover, a_jk’_= a_jk_ +1, and since a_jk_ is smaller than b_jk_, we also have ∣b_jk_ – a_jk’_ ∣ = ∣b_jk_ – a_jk_ ∣ - 1. The only bond we did not choose to change is a_li_ . The order a_ij’_ of the transformed bond is either closer to b_li_ than was an, then the difference of the sum of absolute order differences is reduced by 4, or is further from b_li_, and this sum is reduced by 2. Therefore, if M_a_ and M_b_ are different, we can apply a rule from this set of rules that will decrease the sum of absolute order differences by at least 2 and at most 4.

##### Lemma 3

Considering M_a_ and M_b_ an isomer of M_a_, the 19 rules described in Figure 3 allow us to transform M_a_ into M_b_ using at most 3/4*(*N*^2^ − *N*) single transformations, where *N* is the number of atoms, respecting the following constraints: the maximal valence is 4 and there cannot be two double bonds on the same atom in a 3 or 4 membered ring.

##### Proof

Let us consider M_b_ an isomer of M_a_. If the sum of absolute order differences is not null, then M_b_ is different from M_a_ and using *Lemma* 2, we know we can apply a rule that will strictly decrease the sum of absolute order differences. This sum is obviously positive, is an integer, and is strictly decreasing each time we apply a transformation rule so it will converge to 0 in S/2 transformations at most, where S is the sum of absolute order differences between M_a_ and M_b_. When this sum is null, all bond orders are the same, which means the molecules are the same. An upper estimation of the maximum bond order difference is obtained when M_a_ only has triple bonds, which all have to be deleted. In that case, the sum of absolute order differences is: S = 3*(*N*^2^ − *N*)/2, where *N* is the number of atoms and (*N*^2^ − *N*)/2 the number of defined orders (since a_ij_ = a_ji_). Therefore, since the sum decreases by at least 2, the maximum number of transformations we need to apply is 3*(*N*^2^ − *N*)/4.

#### Proof of the main claim

Given the workings of the algorithm (breadth-first, as explained in section 4.2), the number of iterations for generating all isomers is the number of iterations for generating the furthest one in term of bond order difference from our starting molecule. Therefore, applying Lemma 3, we know the maximum number of iterations of the algorithm is 3*(*N*^2^ − *N*)/4. _

**<u>Notice that</u>** although the number of iterations of the algorithm scales O(*N*^2^), the number of transformation rules applied (i.e.: single reactions) is proportional to the number of isomers.

##### Corollary 1

The maximum number of iterations to generate all alkanes is *N*-1, where *N* is the number of carbon atoms (hydrogens are not considered here).

##### Proof

Adapting the demonstration of Lemma 3, we have to consider the sum of absolute order differences of the farthest isomers that can be reached. Since alkanes are acyclic, the number of bonds is *N*-1 (proven by a simple recurrence, the new atom being joined at a single point to the chain since the molecule is acyclic). Therefore, considering all bonds are different in the new molecule, the sum of absolute order differences is at most 2(*N*-1). Therefore, the maximum number of steps is *N*-1.

The isomer transformation algorithm was applied to generate all alkanes up to 18 carbon atoms using rule *R*_1_ of Figure 3, since it is the only rule with only single bond. Results are presented in Table 1, where it can be seen that *Corollary* 1 is verified in practice.

**Table 1.** Number of generated alkane isomers by canonical augmentation algorithm and isomer transformation algorithm

| Nbr of carbon atoms | Nbr of structures output by canonical augmentation algorithm | Nbr of structures output by isomer transformation algorithm | Nbr of iterations for isomer transformation algorithm |
| --- | --- | --- | --- |
| 1 | 1 | 1 | 1 |
| 2 | 2 | 1 | 1 |
| 3 | 3 | 1 | 1 |
| 4 | 5 | 2 | 2 |
| 5 | 8 | 3 | 3 |
| 6 | 13 | 5 | 3 |
| 7 | 22 | 9 | 4 |
| 8 | 40 | 18 | 5 |
| 9 | 75 | 35 | 5 |
| 10 | 150 | 75 | 6 |
| 11 | 309 | 159 | 7 |
| 12 | 664 | 355 | 7 |
| 13 | 1466 | 802 | 8 |
| 14 | 3324 | 1858 | 9 |
| 15 | 7671 | 4347 | 9 |
| 16 | 18030 | 10359 | 10 |
| 17 | 42924 | 24894 | 10 |
| 18 | 103447 | 60523 | 10 |

### 2.2. Virtual screening in the chemical space

In this section we used RetroPath2.0 to search all molecules that are at predefined distances of a given set of molecules. Such queries are routinely carried out in large chemical databases for drug discovery purposes [31], but in the present case we search similar structures in the entire chemical space. To perform search in the chemical space, we used a source set composed of 158 well-known monomers having a molecular weight up to 200 Da. Our rule set included the transformations colored green in Figure 3 (i.e. transformation rules where double bonds are not transformed into cycles and conversely). For each monomer, RetroPath2.0 was iterated until no new isomers were generated. Each generated structures at a Tanimoto similarity greater than 0.5 from its corresponding monomer were retained (Tanimoto was computed using MACCS keys fingerprints [32]).

Next we wanted to probe if the generated structures exhibited interesting properties as far as polymer properties are concerned. To that end we first developed a QSPR model taking properties from [33]. We focused on polymer glass transition temperature *T_g_* data [34]. The QSPR model was based on a random forest trained using RDKit fingerprints descriptors [19]. The obtained model had a leave-one-out cross-validation performance of Q^2^= 0.75. The model was then applied to predict the *T_g_* for the set of enumerated isomers. Figure 5 compares the distribution of predicted *Tg* values for the enumerated isomers with those obtained from isomer structures available from PubChem. *T_g_* values for enumerated isomers appeared evenly distributed around 301.86±25.69 K compared with the isomers that were available in PubChem (331.66±46.19 K). This shift in the *T_g_* values could be explained by the difference in distribution that necessarily exists between the isomers that are present in PubChem and the total number of enumerated isomers. As we lower the Tanimoto threshold, some monomers might become underrepresented in terms of isomer availability in PubChem. Figure S3 in Supplementary shows the distributions of both sets of isomers in function of the threshold. The increased ability of selecting polymers with *T_g_* above or below room temperature for the enumerated set compared with the PubChem isomers is a desirable feature, as this parameter will determine the mechanical properties of the polymer [35]. In that way, performing a virtual screening of the chemical space of isomers of the reference monomers opens the possibility to engineering applications with improved polymer design.

**Figure 5.**
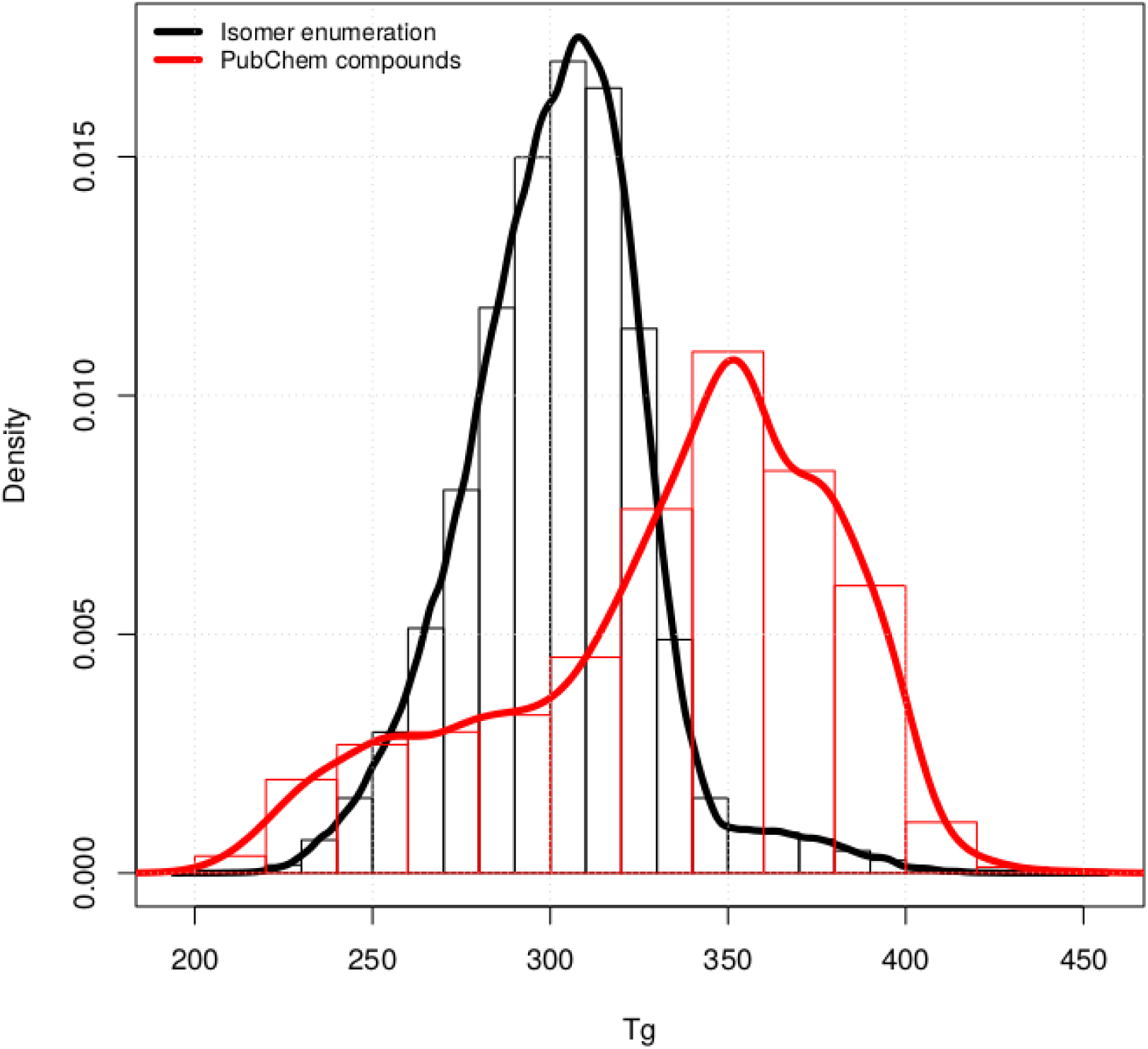
Distributions of predicted T_g_ values for enumerated isomers and for isomers found in PubChem. Distribution of predicted polymer glass transition temperature *T_g_* for enumerated isomers and for isomers found in Pubchem of a reference set of 158 monomers with a Tanimoto similarity greater than 0.5

Moreover, we were interested in determining how many of the starting 158 monomers were accessible through biosynthesis. Namely, how many of the compounds can be synthesized by engineering a metabolic pathway in a chassis organism. This computation can be accomplished by RetroPath2.0 by defining all naturally produced chemicals as sinks in the workflow and using a collection of known enzymatic reaction rules in reversed mode. The process has been described in detailed elsewhere [17]. Through the application of the rules in a retrosynthetic fashion, it is possible to determine the routes that connect the target compounds to the natural precursors. Of the 158 available monomers, using the RetroPath2.0 workflow downloaded from MyExperiment.org [36], we were able to identify 26 compounds that can be naturally synthetized (Figure 6-A). We provide in an archive containing the list of pathways for those 26 compounds.

The QPSR model for *T_g_* was applied to the set of enumerated isomers. As shown in Figure 6-B, the resulting set provided a good covering of the chemical space surrounding the starting monomer set. Moreover, a significant number of enumerated isomers shown a high predicted *T_g_* value, which may indicate a good candidate as a building block replacement for known monomers. Interestingly, those isomers that were close to biosynthetic accessible monomers (Tanimoto based on MACCS keys fingerprint > 0.8) had a distribution of predicted *T_g_* values that significantly differ from the full set (p-value < 1e-12 Welch t-test), with a mean *T_g_* =352.1K (*T_g_* =301.9 K in the full distribution). These close isomers to biosynthetically accessible monomers might be considered as good candidates for alternative biosynthesis since reaching them through biosynthesis may require only few modifications of the original catalytic route.

**Figure 6.**
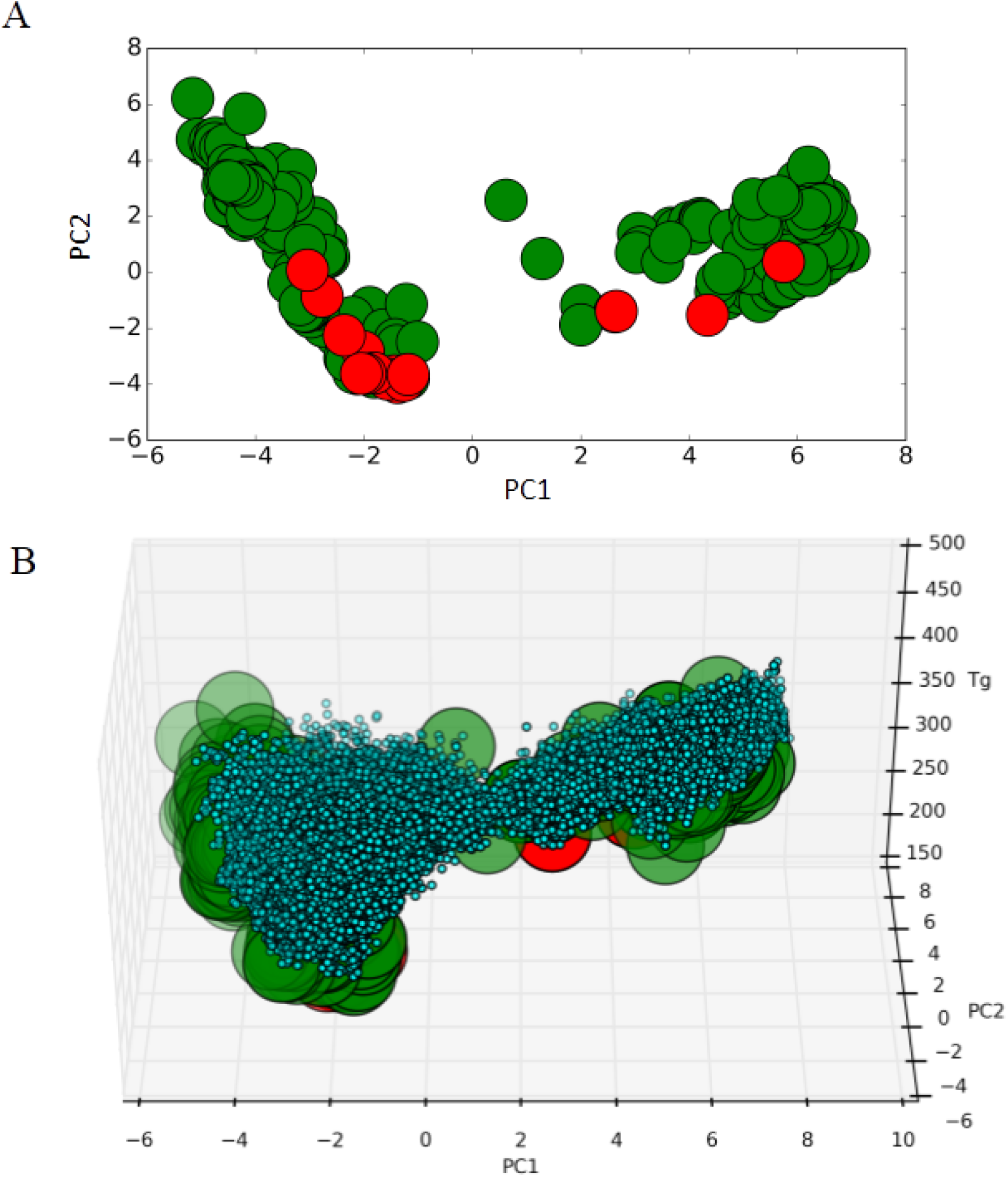
(A) Initial 158 monomers (*green big circles*) represented in the chemical space of chemical descriptors using the two main principal components computed from the MACCS fingerprints as axes. Monomers that can be produced through biosynthesis are represented as big circles in red. (B) Covering of the chemical space generated by the 574,186 isomers (blue) enumerated for the 158 monomers (green) with a Tanimoto similarity greater than 0.5 and associated predicted *T_g_* property of the resulting polymer. Virtual monomers are depicted as small circles to facilitate visualisation of their distribution around the starting monomers.

### 2.3. Search for molecules maximizing biological activities

In this section we are interested in searching chemical structures in the chemical space optimizing biological activities. This type of search can be solved using inverse QSAR procedures [37]. Inverse QSAR requires to first building a QSAR equation predicting activities from structure and then either (i) inverting the equation and enumerating structures matching a given activity [37] or (ii) searching in the chemical space structures similar to those used to build the QSAR equation [33] but having optimized activities. The second approach makes use of either deterministic methods such as lattice enumeration [38] or stochastic searches.

We propose here to use RetroPath2.0 to solve the inverse QSAR problem using a stochastic approach with isomer transformation rules and enzymatic rules for biosynthetic accessibility. To this end, we selected a dataset of 47 aminoglycosides structures for which antibacterial activities have been measured using a MIC assay [39]. The dataset is composed of natural aminoglycosides (gentamicin, tobramycin, neomycin, kanamycin A and B, paromomycin, ribostamycin and neamine) to which are added synthetic structures build on a neamine scaffold. This dataset has already been used to build a QSAR model based on CoMFA analysis leading to a Q2 of 0.6 for a Leave-One-Out (LOO) procedure [39]. We provide in Supplementary a QSAR workflow that makes use of RDKit fingerprints [19] and random forest as a learner leading to a higher Q2 (0.7) for LOO. With that QSAR in hand we run RetroPath2.0 with a source set composed of the 47 aminoglycosides used in the training set, and two different reaction rules sets. The first set is extracted from the transforming enumeration rules depicted in Figure 3, the second set is composed of enzymatic reaction rules leading to neamine (an aminoglycoside) biosynthesis from glucose. Reaction rules for the second set were computed as explained in the method section resulting in 94 rules specific to the biosynthesis of aminoglycosides.

In both cases, reactions rules were fired on the initial source set composed of 47 structures. All rule products were ranked according to their predicted activities as calculated by the QSAR and were selected for the next iteration according to a tournament procedure describe in the method section which derives from [40]. The Figure 7 (A and B) below gives the top activity and the average population activity vs. iteration. The most active structures found by each rule set are also drawn in Figure 7 (C and D).

**Figure 7.**
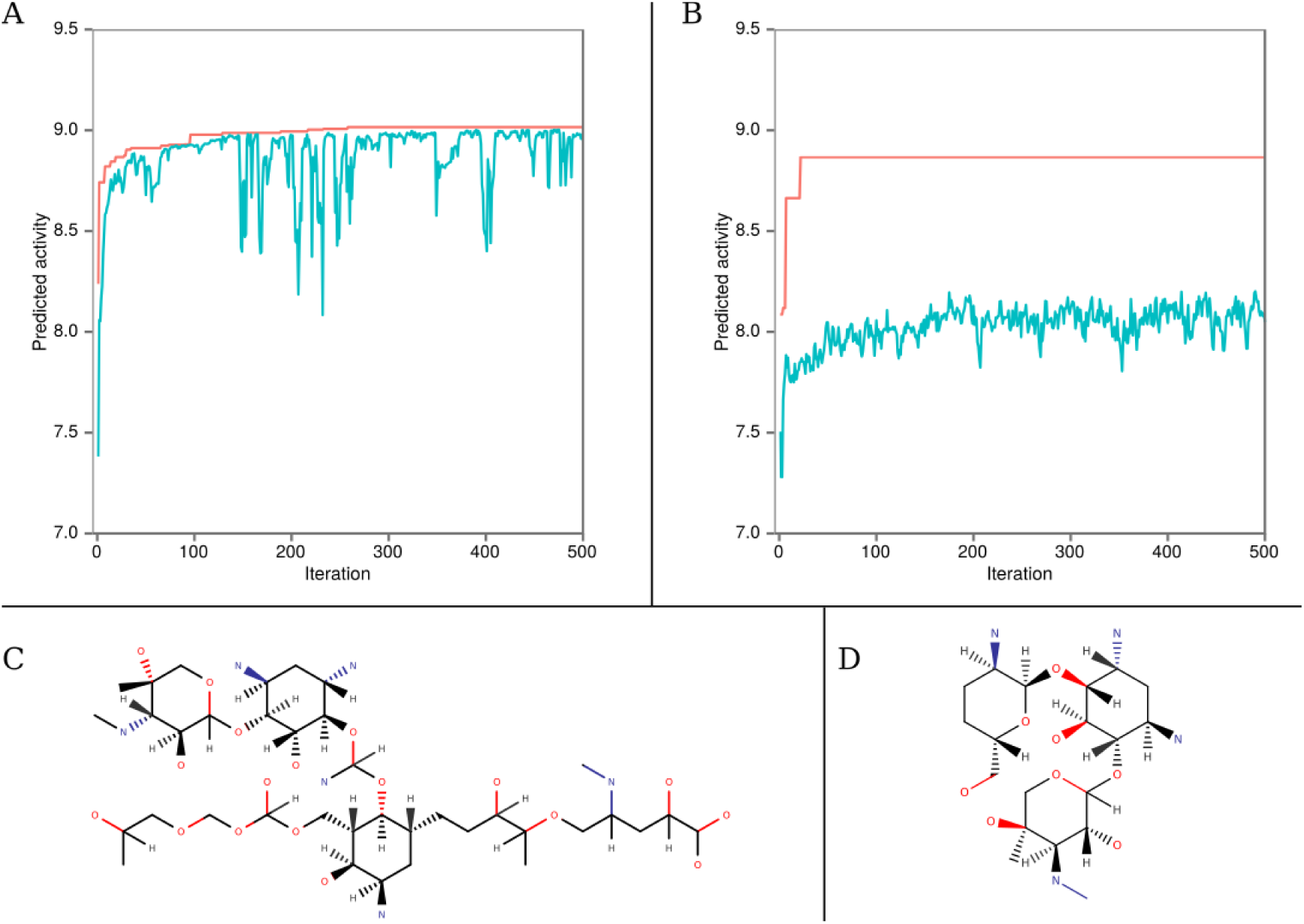
(A) and (B) Evolution vs. iteration number of the best predicted activity (red) and average population predicted activity (blue) from amongst the newly generated structures using (A) transformation enumeration rules or (B) enzymatic rules. (C) and (D) Selected best structure generated after 500 iterations using either (C) transformation enumeration rules or (D) enzymatic rules.

We observe in Figure 7 that the average curve in B is lower to the one in A. This is due to the fact that enzymatic rules generated a lot of compounds that are structurally far from aminoglycoside (i.e.H_2_O, NH4^+^, O_2_…). Moreover, the rules used for A, allows more transformation / modification, thus enabling to better explore the chemical space, and ultimately finding more active compounds. We note that the structure in Figure 7 C have a slightly better predicted activity (pACT = 9.015) than the initial compounds used in the training set, while the structure in Figure 7 D have the same predicted activity than gentamicin (pACT = 8.867).

### 2.4. Metabolome completion and metabolomics

In this last example we use enzymatic reaction rules in an attempt to complete the metabolome of species used in biotechnology. We are motivated here by current efforts invested to complete the knowledge on the metabolism of various organisms [6, 7, 15]. The benefits are numerous and include the identification of relevant biomarkers for many diseases; for personalized nutrition advice; and also for searching for relevant indicators and metabolites of plant and animal stress in agricultural practices and breeding programs. Additionally, knowing the metabolic space of microbes is an essential step for optimizing metabolic engineering and creating synthesis pathways for new compounds for industrial applications.

Experimental evidences from metabolomics analyses are often informing us that with currently known metabolites one cannot cover the ranges of masses found in actual samples, and consequently there is a need of completing the metabolomes of interest. This need is clearly seen in the Human Metabolome Database (HMDB) where the number of reported masses has recently grown from 20,931 in 2013 [41] to 74,461 (at the time this manuscript was written), while annotated metabolites in metabolic databases are still in the range of 1847 (HumanCyc). Despite such a growth in databases, a significant amount of spectral peaks remains unassigned. This high fraction of unassigned peaks might be due to several factors including isotope, adduct formation, ion fragmentation, and multimers. Besides such sources of uncertainty in samples, many unassigned peaks should also be due to promiscuous activities of enzymes not yet characterized because of the lack of an appropriate description of the mechanisms of enzyme promiscuity.

To gain insights into those mechanisms enabling promiscuity, reaction rules have been shown to be appropriate [21] in particular the rules allowing to focus on the center of the reactions. To this end, several enzymatic reaction rules have been proposed such as those derived from bond-electron matrices [42], on the smallest molecular substructure changing during transformations [9], or on reaction rules that codes for variable environments at reaction centers (see [7] and method section). That latter reaction rule system codes for changes in atom bonding environments where the reaction is taking place and the environment can range from including only the atoms participating to the reaction center to the entire set of atoms and molecules participating to the reaction. The advantage of that latter approach is that the size of the environment (named diameter) can be tuned to control the combinatorial explosion of possible products.

The degree of plasticity in metabolic networks that is uncovered by variable reaction center diameter is actually revealing an intrinsic feature of organisms linked to their adaptability, i.e. enzyme promiscuity. Promiscuity stands for the ability of enzymes to catalyze more than one reaction or to accept more than one substrate, a mechanism, which can be traced to the evolutionary origins of enzymatic functions. Mimicking nature, such enzyme versatility can provide novel ways for biosynthesizing metabolite and even bioproducing non-natural molecule. To that end, the variable diameter method has shown itself to be specially well-suited for modeling the mechanisms of enzyme promiscuity as it has already enabled the experimentally validated discovery of a novel metabolite in *E. coli* and of the promiscuous enzymes producing it [21].

In this study, we make use of RetroPath2.0 to exemplify how variable reaction rule diameters can be used to complete the metabolome of *E. coli.* More precisely, we used as a source set all the metabolites present in *E. coli* iJO1366 model [43]. We first tested two rules sets aforementioned, a set of about 100 reaction rules part of the BNICE framework [42] and a set of 50 reaction rules developed with the Sympheny software [9]. The reaction rules coded in the form of SMARTS string are provided in Supplementary at MyExperiment.org, along with the EC numbers corresponding the rules. While the two rule sets were not developed to code only for *E. coli* reactions, for each EC number there is a corresponding enzyme annotated in *E. coli* so we kept all rules in the two sets. We then tested reaction rules with variable diameters using the procedure described in the method section to code for all *E. coli* metabolic reactions extracted from iJO1366 model. Rules were calculated for each reaction with diameters ranging from 2 to 16. The Table below provides the number of compound generated running RetroPath2.0 for one iteration on the metabolites of the iJO1366 model and the rules sets mentioned above (see Method section for additional details).

Table 2 shows that the number of compound generated increases as the diameter decreases. This is consistent with the fact that shorter diameters will accept more substrates than higher ones and will thus produce more products. Although they were not constructed with diameters, the BNICE and Sympheny rule sets generally correspond to small environments comprising only few atoms and bonds around reaction centers, which explain why these two systems generate more products than high diameter rule sets. Nonetheless, even with high diameters, all variable diameter rule sets produce more molecules found in *E coli* model than the BNICE and Sympheny rule sets. This might indicates that the variable rule sets correspond to a more accurate coding of metabolic reactions than the other systems.

**Table 2.** Compounds generated by RetroPath2.0 using various reaction rules applied on *E. coli* iJO1366 model metabolites [43]. All numbers correspond to compounds having different InChIs at the connectivity level.

| Reaction rule set | Nbr compounds generated | *E. coli* model coverage (1) | MS peaks coverage (2) | Median nbr cmpds per peak | Averaged nbr cmpds per peak |
| --- | --- | --- | --- | --- | --- |
| *E. coli* Model [43] | 751 | 100.0 | 12.3 | 1 | 1.5 |
| Sympheny [9] | 9448 | 48.2 | 40.4 | 3 | 6.3 |
| BNICE [42] | 8421 | 68.8 | 45.3 | 3 | 5.8 |
| D16 (3) | 1230 | 82.0 | 23.1 | 1 | 2.0 |
| D10 | 2992 | 83.6 | 25.6 | 1 | 2.3 |
| D8 | 5055 | 84.2 | 28.1 | 1 | 2.7 |
| D6 | 11981 | 84.7 | 46.6 | 2 | 3.1 |
| D4 | 37450 | 86.8 | 60.6 | 2 | 5.7 |
| D2 | 162480 | 91.7 | 79.9 | 8 | 16.9 |

(1) The *E. coli* model contains 751 compounds (with different connectivity InChIs). The column report the % of these 751 compounds generated by the different rule sets. (2) The MS spectra were downloaded from Metabolight [44] and the OpenMS workflow described in the Method section retrieved a total of 800 distinct peaks. The column report the % of peak assigned to at least one compound generated by the rule sets. (3) The number indicates the diameter

To further probe the coverage of the various rules sets listed in Table 2 we searched if the compounds produced could be found in MS spectra. To this end, we downloaded MS spectra from Metabolight [44] where masses have been measured on *E. coli* cell extracts. The spectra downloaded corresponded to a study aimed at probing the dynamics of isotopically labeled molecules (i.e. ^13^C labeled glucose) [45]. Since we are concerned here with wild type *E. coli* metabolome, we considered only the spectra where *E. coli* cells had not yet been exposed to labeled glucose (spectra acquired at time t=0). All compounds generated by our various rules sets were prepared to be read by OpenMS nodes [23] and a workflow was written with these nodes to annotate the MS spectra peaks (cf. Method section for details).

The results presented in Table 2 shows that as the diameter decreases the number of peak assignment increases, which is not surprising considering that the number of compounds generated increases as well. We observe that the Sympheny and BNICE rules sets give results similar to those obtained by the D6 rule set, albeit with a higher number of annotations per peak.

In all cases the rules sets produced compounds not present in the *E. coli* model but with corresponding masses in the MS spectra. Table S2 in Supplementary give a list of 40 such compounds having an identifier in MetaNetX [11] and produced by three identical reactions (i.e., reactions having the same substrates and products) generated using the Sympheny, BNICE and D6 rule sets. The compounds were produced by 53 reactions, some compounds being produced by more than one reaction. We note that the 40 compounds have been generated by rule sets for which at least one gene in *E. coli* has been annotated with the same corresponding EC number. The 40 compounds are thus potential new *E. coli* metabolites and their presence should be further verified using for instance MS/MS analysis.

### 3. Conclusions

In this paper we have presented a general method allowing one to explore the chemical space around a given molecule, or around a given set of molecules. The originality of the method is that the exploration is performed through chemical reactions rules. We have given a set of rules allowing us to generate any isomer of any given molecule of the chemical space. We also provide examples making use of reaction rules computed from enzymatic reactions. Using rules computed on known reactions has a definite advantage regarding the (bio)synthetic accessibility of the molecule produced, which not necessarily is the case for other techniques producing molecules *de novo* [33,37,40,49,50,51,52].

Our method has been implemented into RetroPath2.0, a workflow running on the KNIME analytics platform [18]. RetroPath2.0 can easily be used with source molecules and reaction rules different that those presented in the paper. For instance the workflows provided in Supplementary can be used with the reaction SMARTS rules and fragment libraries (as source compounds) of the DOGS software (inSili.com LLC [4]) developed for *de novo* drug design, other technique evolving molecules toward specific activities or properties [40, 50-52] could also be implemented in RetroPath2.0 provided that one first codes reaction rules in SMARTS format.

Aside from searching molecules having interesting properties and activities RetroPath2.0 can also be used to complete metabolic maps by proposing new metabolites biosynthesized through promiscuous enzymes, these new metabolites can in turn be used to annotate MS spectra and to that end we provide an interface with OpenMS [23]. Finally, RetroPath2.0 was originally developed to enumerate pathways producing a given target product from a source set of reactants. While we have benchmarked the workflow in the context of metabolic engineering [17] it can also be used for synthesis planning as long as synthesis reaction rules are available.

### 4. Methods

#### 4.1. Generating reaction rules

All our reaction rules are represented in the form of reaction SMARTS [14]. Reaction rules used for canonical augmentation are provided in Figure 1 and for isomer transformation in Figures 3, 4 and S2. Enzymatic reaction rules were computed taking enzymatic reactions from MetaNetX version 2.0 [11]. To compute rules, we first performed an Atom-Atom Mapping (AAM) using the tool developed by [46] (Figure 8 A). Next, multiple substrates reactions were decomposed into components (panel C and D in Figure 8). There are as many components as there are substrates and each component gives the transformation between one substrate and the products. Each product must contain at least one atom from the substrate according to the AAM. This strategy enforces that only one substrate can differ at a time from the substrates of the reference reaction when applying the rule.

The following step consisted in computing reactions rules as reaction SMARTS for each component. We did it for diameters 2 to 16 around the reaction centre (panels C and D in Figure 8) by removing from the reaction components all atoms that were not in the spheres around the reaction centre atoms.

We extracted more than 24,000 reaction components from MetaNetX reactions, each one of those leading to a rule at each diameter (from 2 to 16).

We provide in Supplementary at MyExperiment.org a subset of 14,300 rules for *E. coli* metabolism. The rules were selected based on the MetaNetX binding to external databases and the iJO1366 whole-cell *E. coli* metabolic model [43]. We also provide enzymatic rules enabling the biosynthesis of aminoglycosides from Glucose. The reactions were extracted from the map00524 KEGG map [47], and rules were computed as above on reactions for which a MetaNetX identifier could be retrieved. The resulting set comprised 94 rules calculated for each diameters ranging from 2 to 16.

**Figure 8.**
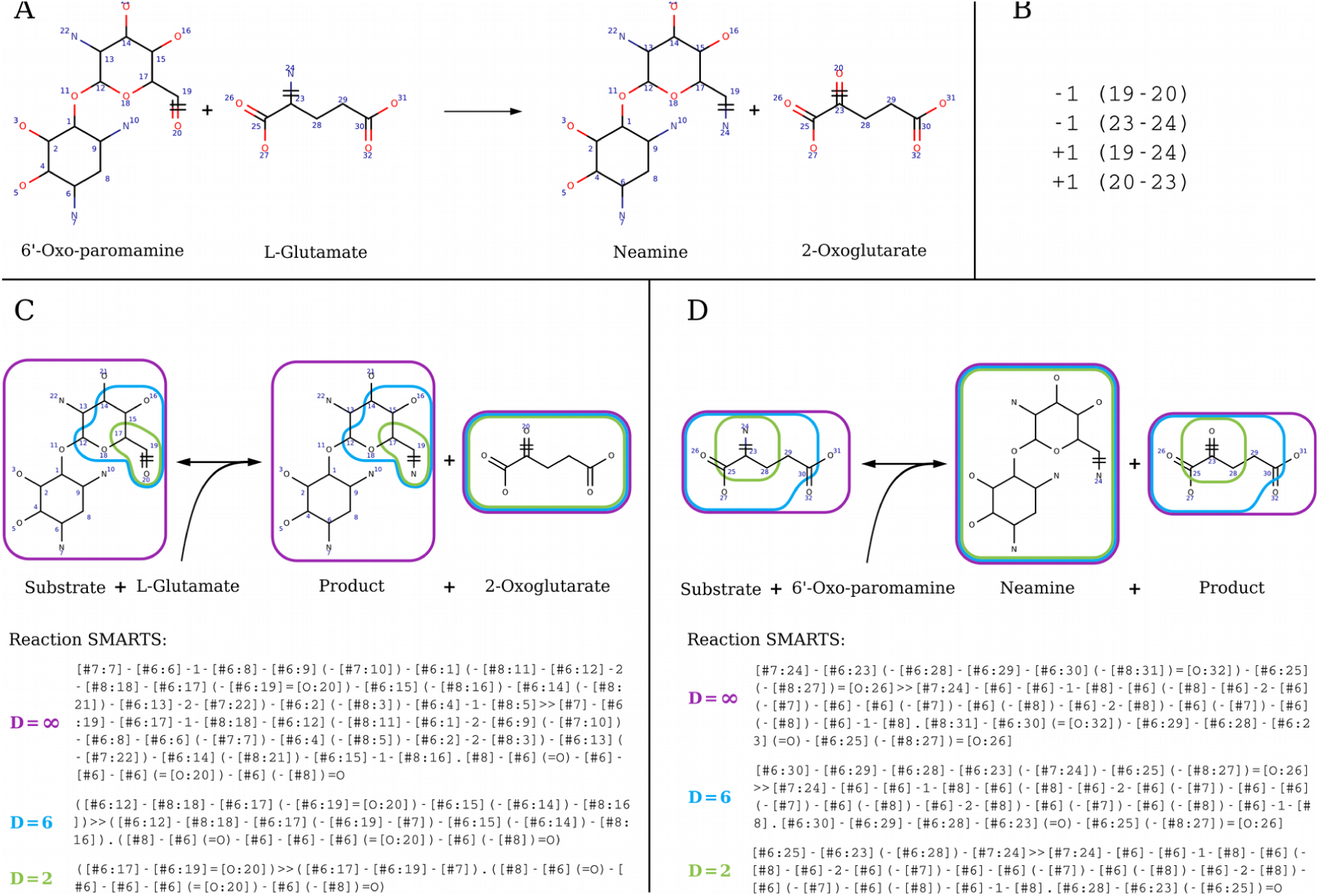
RetroPath2.0 rules and corresponding SMARTS for reaction 2.6.1.93 at various diameters. (A) Full reaction 2.6.1.93 with atom mapping. (B) The list of broken bonds (-1) and bonds formed (+1) is given by their atom numbers. (C) The corresponding SMARTS for the component modelling promiscuity on 6’-Oxo-paromamine: Substrate + L-Glutamate = Product + 2-Oxoglutarate. (D) The corresponding SMARTS for the component modelling promiscuity on L-Glutamate: Substrate + 6’-Oxo-paramamine = Neamine + Product. C and D. Rules are encoded as reaction SMARTS and characterized by their diameter (∞ purple, 6 blue or 2 green), that is the number of bonds around the reaction centre (atoms 19, 20 and 23, 24) defining the atoms kept in the rule.

#### 4.2. RetroPath2.0 core algorithm

The RetroPath2.0 workflow essentially follows an algorithm proposed by some of us [16, 17] and its workflow implementation, which has already been described in details [17], is summarized in Figure 9. We here focus on the different usages of RetroPath2.0 for the use cases provided in section 2.

**Figure 9.**
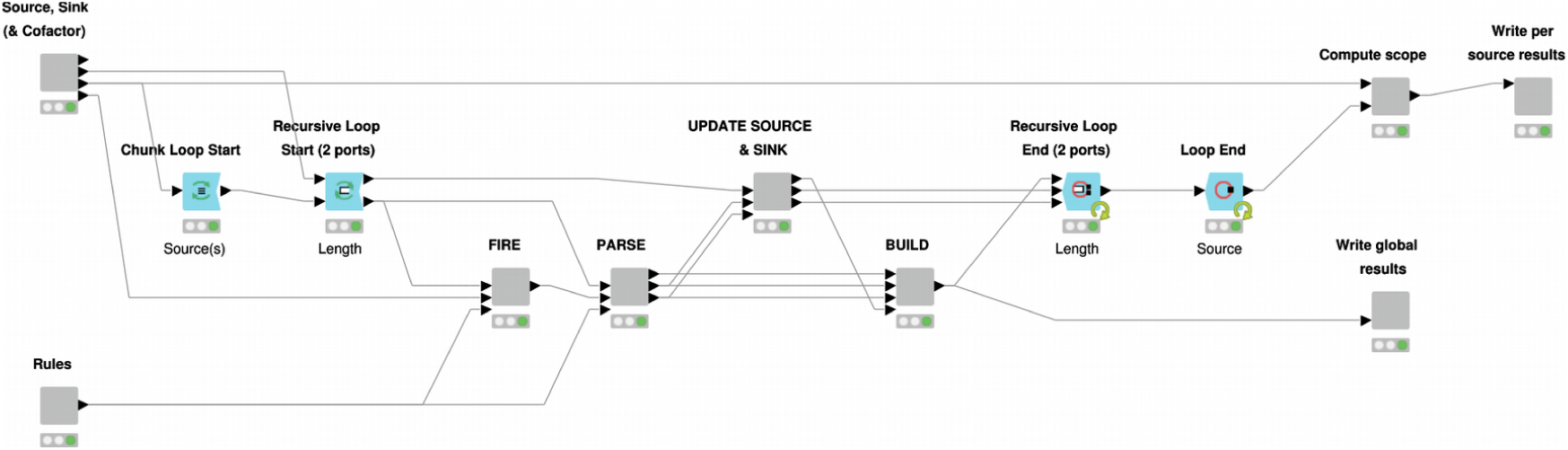
RetroPath2.0 KNIME workflow. Inner view of the “Core” node where the computation takes place. The “Source, Sink…” and “Rules” nodes parse the source, sink and rules input files provided by the user and standardize data so that it can be processed by downstream nodes. Definitions for source, sink, and rule sets are provided in the text. The outer loop (“Source” loop) iterates over each source compounds, while the inner loop (“Length” loop) allows to iterate the process up to a maximum number of steps predefined by the user. The nodes (i) “FIRE”, (ii) “PARSE”, (iii) “UPDATE SOURCE…” and (iv) “BUILD” are sequentially executed at each inner iteration. Respectively, they (i) apply all the rules on source compounds, (ii) parse and standardize new products, (iii) update the lists of source and sink compounds for the next iteration and (iv) merge results that will be written by the node “Write global results”. Once the maximum number of steps is reached (or no new product is found), the “Compute scope” node identify the scope linking each source to the sink compounds, then these results are written by the node “Write per source results”. Only the main nodes involved in the process are shown.

In all cases the workflow performs the generation of structures in a breadth-first way by applying iteratively the same procedure. An iteration starts by applying reaction rules to each of the compounds of a source set. For each compound, the products are computed using the RDKit KNIME nodes one-component or two-component reactions [19]. Products are sanitized (removal of structures having incorrect valence), standardised and duplicates are merged. The set of products will become the new source set for the next iteration. The workflow iterates until a predefined number of iterations is reached or until the source set is empty.

In the case of isomer augmentation (workflow RetroPath2.0-Mods-isomer-augmentation, sections 2.1) the initial source set is composed of a single carbon atom and the rule used is *R*_1_ in Figure 1, since it is the only rule that will produce acyclic molecules. The rule is fired on the source set, and the products become the new source set in the next iteration. The workflow is iterated a number of times equal to *N*-1, where *N* is the number of atoms one wishes the final molecule to have.

In the case of isomer transformation (workflow RetroPath2.0-Mods-isomer-transformation, sections 2.1 and 2.2) the initial source set is composed of a molecule that is filled with the appropriate number of hydrogens using the RDKit KNIME node Add Hs. At each iteration rules are fired on the source set and the products obtained become the new source set for the next iteration. As an additional last step of each iteration, products that have already been processed in a previous iteration are filtered out before building the next source set. This necessitates maintaining a set (named sink) comprising all molecules so far generated. All products that have already been obtained are removed from the product set and the remaining molecules are (i) added to the sink set and (ii) used as the new source set for the next iteration. This avoids applying reactions on the same products during subsequent iterations. Disconnected structures are removed from the results by filtering out any product having several disconnected components (according to the SMILES representation). When enumerating alkane, disconnected structures represents between 50 and 66% (depending of the alkane size) of the generated structures before filtering and merging duplicates. To generate the results of Table 1, since we are enumerating alkanes (no multiple bonds or cycles), the rule to be used is *R*_1_ in Figure 3. To enumerate the isomers of the monomers in section 2.2, if we prohibit the transformation of multiple bonds into cycles and thus keep the number of single, double and triple bonds constant, the rules to be used are *R*_1_, *R*_5_ and *R*_9_ in Figure 3 (also found in Figure S2 since the monomers used do not contain triple bonds). Since this algorithm can become computationally intensive, we also provide an additional workflow (called RetroPath2.0-Mods-isomer-transformation-queue) to deal with memory management. This workflow illustrates how to introduce a FIFO data structure for the source set (i.e. queue containing structures upon which rules will be fired) and use it for iteratively firing rules on small chunks of structures (e.g. chunk of 20 structures), new products obtained are then added to the source queue. Interestingly, the breadth-first approach for generating the structures can be replaced by a depth-first approach by replacing the queue (first in, first out structure) by a stack (last in, first out structure).”

In the case of inverse-QSAR (workflow RetroPath2.0-Mods-iQSAR, section 2.3), the source set initially comprises the molecules used in the training set when building the QSAR. At each iteration, one or two molecules are chosen at random from the source set depending on the rule set that is being used (one molecule with enzymatic reaction rules, two molecules with isomer transformation rules). Rules are then fired on the selected molecules and an activity is predicted for each product using the QSAR equation. The source set is updated retaining molecules according to a selection tournament procedure borrowed from [40]. Briefly, the initial source set (i.e. the set of structures used at the start of the current iteration) is merged with the product set (i.e. the set of structures obtained after firing the rules). This merged set is then randomly split into 10 subsets and the 10 top best structures from each subset are retained according to their predicted activity. Finally, all the retained structures are pooled together to form the updated source set to be used at the next iteration. The workflow is iterated a (user) predefined number of times.

In the case of *E. coli* metabolic network completion (workflow RetroPath2.0-Mods-metabolomics, section 2.4), we provide three workflows. The first workflow is RetroPath2.0, which is fully described in [17] and is similar to the isomer transformation one. Here, RetroPath2.0 produces a list of molecules obtained using *E. coli* enzymatic reaction rules (see section 3.1). The second workflow takes as input the products generated by RetroPath2.0, computes the exact mass for each product and prepare files to be read by OpenMS nodes for MS data peak assignment [23]. The last workflow is build with OpenMS nodes, it reads several MS data file in mzML format, two lists of adducts in positive and negative modes, and the files generated by the second workflow (containing RetroPath2.0 generated product with masses). The workflow searches for each compound the corresponding peak in the MS spectra. The workflow was parameterized for metabolomics analysis as described in OpenMS manual [48], the AccurateMassSearch node was set to negative ion mode as the experiment were carried out with an LTQ-Orbitrap instrument operating in negative FT mode (cf. protocols in [44]).

Further details on how to run all the above workflows are provided in the Supplementary at MyExperiment.org

## Authors’ contributions

The work was directed by JLF who sketched the proof of the claim, designed all workflows, and generated the results presented in sections 2.1 and 2.4. MK wrote the proofs in section 2.1, TD wrote all workflows and produced the results in section 2.3. PC used the workflows to generate the results in section 2.2. All authors contributed to the manuscript write-up.

## Acknowledgements

The authors acknowledge Baudoin Délépine for his contribution to the general workflow architecture and enzymatic reaction rules calculations.

## Availability of data and material

The workflows and associated data supporting the conclusions of this article are available on MyExperiment repository, http://www.myexperiment.org/packs/728.html.

## Ethics approval and consent to participate

N/A.

## Consent for publication

N/A.

